# A modeling of a Diphtheria epidemic in the refugees camps

**DOI:** 10.1101/208835

**Authors:** María Torrea, José Luis Torrea, Daniel Ortega

**Author notes:** ∗*Corresponding author: Servicio de Urgencia, Hospital General Universitario Gregorio Marañón, 28007 Madrid, Spain.

## Abstract

**Background:** Diphtheria has a big mortality rate. Vaccination practically eradicated it in industrialized countries. A decrease in vaccine coverage and public health deterioration cause a reemergence in the Soviet Union in 1990. These circumstances seem to be being reproduced in refugee camps with a potential risk of new outbreak.

**Methods:** We constructed a mathematical model that describes the evolution of the Soviet Union epidemic outbreak. We use it to evaluate how the epidemic would be modified by changing the rate of vaccination, and improving public health conditions.

**Results:** We observe that a small decrease of 15% in vaccine coverage, translates an ascent of 47% in infected people. A coverage increase of 15% and 25% decreases a 44% and 66% respectively of infected people. Just improving health care measures a 5%, infected people decreases a 11.31%. Combining high coverage with public health measures produces a bigger reduction in the amount of infected people compare to amelioration of coverage rate or health measures alone.

**Conclusions:** Our model estimates the evolution of a diphtheria epidemic outbreak. Small increases in vaccination rates and in public health measures can translate into large differences in the evolution of a possible epidemic. These estimates can be helpful in socioeconomic instability, to prevent and control a disease spread.

## 1 Introduction

Diphtheria is a highly contagious disease with a big mortality rate especially in children. Vaccination campaigns have made it practically eradicated in industrialized countries. In Spain, without any case since 1986, see [1], the last case registered was in 2015 in Olot in a non vaccinated child. However, if the vaccination coverage were to fall, the disease could emerge again. This statement is backed by the epidemic outbreak in the Soviet Union in 1990. This epidemic has been the object of several researches and discussions, [2], [3], [4], [5]. There is a common land of coincidence of these researches. In one hand the decrease of vaccination coverage, essentially due to a great economic, social and political crisis. On the other hand, the crisis also pushed down the public health care. These two vectors together with the impoverishment, overcrowding and movements of large population groups propitiated the breeding ground for the above-mentioned epidemic outbreak, [6].

In this article we are interested in a possible Diphtheria epidemic among “refugees”. At present, under the generic name of refugees are designated different movements of population around the world, mainly in the Mediterranean area. All of them with conditions of extreme overcrowding, lack of hygiene and basic health care. Ve shall focus in the groups coming from Syria. The Office of the United Nations High Commissioner for Refugees (UNHCR) estimate in more than 5 million internal syrian refugees between Turkey, Jordan, Egypt and Greece, [7]. In our opinion, the pattern of the Soviet population of the 1990's is being reproduced almost exactly. Namely, refugees are a well sanitized population until a few months ago, even with the immunization record in order, but they are in a situation of total economic and social deterioration. This leads to a decrease in vaccination surveillance (despite international emergency care protocols), as well (and no less important) to a shortage of medicines. Our model could be applied to other cohorts like Libyan, Afghans and Sub-saharian.

In this article we construct a mathematical model that describes the evolution of the outbreak epidemic in the Soviet Union. Once the mathematical model is developed, we can ask ourselves questions about the epidemic evolution like the following:

- How could have been its evolution with a better vaccine coverage?
- How about with a better public health care?
- How about a combination of a better public health care together with a small increase of vaccine coverage?
- And even with a worst vaccine coverage?

These questions are implemented in our model giving us outputs with quantitative numbers and illustrative graphs and tables. The purpose of this article is to applied these outputs to the case of refugees. In our opinion, it is difficult to change the conditions of deteriorated human life in the refugees camps. The conditions of overcrowded and big movements of population seem to be, sadly in Europe, one of the human disasters of our days. However a small increase of health care could be possible. In particular a surveillance of potential patients, a quick treatment protocoled measures, see [8], [9], and a (possible) small increase of vaccination. All these measures would produce a big reduction in the number of infected people. What is more, controlling the epidemic is not only important for the places where the refugees are, since some cases of cutaneous diphtheria have already been reported in Denmark, Germany and Sweden, see [10], [11]

We present our results in Section 3. In section 4 are exposed our conclusions. Figure 7 in Section 4 where epidemic is evaluated in differents scenarios of vaccine coverage and health measures is the graphic summary of this article

## 2 Material and methods

### 2.1 Illness description

Diphtheria (from the Greek *διϕθερα* “membrane”) is caused by Corynebacterium diphtheriae, a gram positive bacillus, whose main virulence factor is the exotoxin produced by some strains that cause local and remote cell destruction. Non-toxigenic strains produce a symptomatic, generally milder, disease ([9], [12]). Four biotypes have been identified: gravis, mitis, intermedius and belfanti. This is important for the posibility that changes in the circulating strains of C. diphtheriae could be responsible for cyclicality and episodic epidemic waves associated with the incidence of diphtheria in the pre-vaccine era ([13]). In most cases, a carrier state occurs as an asymptomatic oropharyngeal level. However, in severe cases it produces an adherent greyish pseudomembrane that affects the upper respiratory and digestive tract and may cause airway obstruction. In addition, its generalized toxigenicity is associated with complications in other organs (myocardium, kidneys and central nervous system) with a lethality rate of more than 10% ([14], [15]).

The incubation period varies from 1 to 7 days. Humans are the only reservoir. The transmission is by contact of nose, throat, eyes and skin secretions from infected people. Rarely from contaminated food or tools. Infectivity from an untreated patient usually last for 2 weeks, but may persist several months. In adequately treated patients infectivity lasts less than 4 days ([8]). Vaccinated subjects can be carriers of the disease and also contagious. In endemic areas up to 3.5% of the population becomes a carrier, while in countries with a current vaccination schedule the isolation of the microorganism in healthy patients is extremely rare ([15]). Incidence is high in autumn and winter and mostly affects low socioeconomic groups living in overcrowding and with limited access to health services. If a case is suspected, the patient should be isolated and the treatment started immediately after taking the bacteriological samples, without waiting for laboratory confirmation. Diphtheria antitoxin is the key element in the treatment and should be given as soon as the disease is suspected, as it improves prognosis. Patients should also receive antibiotics to remove the bacteria, the duration of the transmissibility period and carrier status. Treatment should be continued for 14 days. It is also recommended to initiate or complete active immunization (vaccine) during the convalescent period, as the disease does not always confer immunity ([8], [16]).

Fortunately diphtheria is prevented by vaccination against diphtheria, tetanus and acellular pertussis (DTPa / Tdpa). The Vaccine Advisory Committee of the Spanish Association of Pediatrics (ACV-AEP) in 2017 recommends 5 doses: first vaccination with two doses (2 and 4 months) of DTPa (hexavalent); 12-month (3rd dose) booster with DTPa (hexavalent) at 6 years (4th dose) with standard loading preparation (DTPa-IPV) preferable to low antigenic loading of diphtheria and pertussis (Tdpa-IPV) and at 11-14 years (5th dose) with Tdpa [17]. Nowdays in Occidental world, diphtheria is seen as an essentially disappeared disease. Even medical population has no practical experience with it. As a possible canonical example we present the following data relative to Spain. Vaccination covers 90-95% of children and the effectiveness of the vaccine is estimated at 97%. Since 1986 there has been no case in Spain except the one registered in Olot in 2015. However, due to its severity and contagiousness is a notifiable disease in Spain. In the last decades of the nineteenth century and first of the twentieth was the leading cause of child mortality in industrialized countries and in Spain represented a total of 80,879 deaths between 1880 and 1885 ([18]). Throughout history, diphtheria, has produced devastating outbreaks. During the large diphtheria epidemic that occurred in Europe and the United States in the 1880s, case fatality rates of up to 50% were reached in some areas. During World War I, fatality rates declined in Europe around 15%, mainly due to the common treatment with antitoxins. In the World War II, Europe was also affected by diphtheria epidemic, which caused around 1 million cases and 50,000 deaths in 1943. During the 1940s and 1950s, the introduction of universal child immunization with diphtheria toxoid almost eliminated diphtheria in most of industrialized countries. In developing countries, high levels of child vaccinations with three doses of diphtheria-tetanus-pertussis (DTP) vaccine were achieved following the implementation of the World Health Organization's Expanded Program on Immunization (WHO) in the 1970s ([19]). During the vaccination era, anti-vaccination groups started to develop. Despite their arguments ([20], [21]), in our opinion their attitude and philosophy was very dangerous for themselves (the Olot boy who died of diphtheria in 2015 was not vaccinated) as well as for the rest of the population, since it can create a serious public health problem.

Nowadays in immunized populations, diphtheria cases are isolated and limited to family or community groups. However in many underdeveloped countries it remains a public health problem, especially in Asia (in particular India, Nepal and Bangladesh), Southeast Asia, the Pacific (Sub-Saharan Africa), South America (Brazil), and the Middle East (Iraq and Afghanistan), although in recent years the use of vaccine is increasing. In 2007, 15 countries in Asia, Africa, the Middle East, the Caribbean and Europe reported 10 or more cases of diphtheria to the World Health Organization, with a total of 4,190 cases registered worldwide in that year. It is noteworthy that more than 3,000 cases were reported in India, where diphtheria remains endemic and there is documented evidence of lack of immunization as a major risk factor. In 2011, of 4,880 cases reported, 3,485 were from India ([22]). In South America, WHO's Expanded Program on Immunization achieved a drastic and sustained decline in the incidence of diphtheria ([14]).

### 2.2 Diphtheria in the former URSS

In spite of data above, during the 1980s and early 1990s there were small diphtheria epidemics in some countries (Sweden, Germany, Portugal). As possible reasons, adults may have been vaccinated but protection had declined due to lack of memory and not to be in contact with the bacteria owing to the practical disease eradication. Some studies estimates in 19 years the immunization ([24]). These outbreaks did not refiect any public malpractice of vaccination policy. However in 1990 a real epidemic emerged in the countries of the former Soviet Union ([23]). Adults were mainly affected. The epidemic appeared in the countries of the Russian Federation in 1990, but spread rapidly during 1991-1993 by the so-called New Independent States and reached its maximum incidence in 1994-1995 with an annual average of 17 cases / 100,000 habitants and in some areas such as Tajikistan to peaks of 73 cases / 100,000. From 1990 to 1998 more than 157,000 cases and 5,000 deaths were registered, representing 80% of cases of diphtheria registered in the world. See the attached table at the end of this section. In order to understand the reasons for this health emergency, it is necessary to take a historical tour on the disease evolution in the former USSR. Although a vaccination program began in the late 1920s, it was not until after the World War II that we can speak about massive child immunization programs. With some ups and downs, the epidemic declined by more than 90 % in 1963. This downward trend continued and the total of cases registered in the URSS in 1976 were 198 (0.08 / 100,000). A resurgence, mainly in adults, began in the late 1970s, reaching 1,609 cases (0.65 / 100,000) in 1984. Special immunization campaigns succeeded in reducing these numbers and by the end of 1989 the situation was considered under control with 0.34 / 100,000 cases. This earlier optimistic moment produced a relaxation of surveillance policies as well as a misleading perception of the eventuality of the epidemic. Moreover, vaccination schedules and even the type of vaccine were changed. Memory doses were spaced out, even suppressed the so-called school-entry booster dose. The risks of the vaccine were questioned. The result of all this maremagnum was a fall to 60-80% in child vaccination.

It should also be taken into account that in 1990 the population aged 40-50 years had not been immunized in their childhood, nor in the special programs developed in the 1980s during the sporadic outbreaks described above. It made this population susceptible. There are several hypotheses about the onset of the epidemic. All of them argue that the decline in health care and large population movements were the perfect breeding ground for the situation to explode. Refugees movements from Afghanistan to Tajikistan, mass displacements of the rural population due to Georgia and Azerbaijan wars, Russian army battalions placed in Afganistan and returned to Moscow as construction battalions (about 100,000 soldiers) were sources for the bacteria so it spread through the major routes of population movements. Even the new recruits were not routinely immunized and became foci of the disease because of their quartering conditions. The disease particularly affected adults and teenagers, revealing that the previous immunization had not been enough, mainly due to lack of memory doses. What is more, there had been a perverse effect. Due to vaccination, the circulation of the bacteria had almost disappeared, so those adults had not been in contact with the bacteria and therefore were deprived of natural immunity.

All this occurred at a time of socio-economic instability, so inevitably it took time to provide an adequate public health response to the health emergency. There were insufficient doses of vaccines, so it was not until 1995 they reached 93% immunization coverage in children as well as 75% in adults. The situation was particularly harsh in some states of the former USSR and in Baltic States with very scarce resources of antibiotics and antitoxins.

The epidemic was particularly disastrous in places of high concentrations of people with low hygiene and high rates of personal contacts. Finally, the deterioration of sanitary structures led to an inadequate transport of immunization doses, losing their effectiveness. Due to the health emergency there was a global mobilization. Different countries sent laboratory kits to countries of the former USSR with less resources, health personnel were trained in logistics, transport, social mobilization, etc. By 1995, many countries began to see the number of infectious diseases declining compared to previous years (1993-94). By the end of 1996 the infection decreased by 60% compared to 1995 (20,215 cases). In 1998, 2,720 cases were registered.

At the beginning of 1999 the largest diphtheria epidemic in the previous 30 years was under control. It had caused more than 157,000 cases and 5,000 deaths. Despite this devastation, some studies estimate that the relative rapid control of the epidemic prevented an additional 560,000 cases and 15,000 deaths, an assertion contained in [2]. The following table can be found in [2].

### 2.3 The mathematical model

We shall use a system of non linear differential equations which describes the dynamic of the epidemic. For each time *t*, we shall divide the population in the following classes. Susceptible, *S*(*t*), people which can be infected. Infected, *I*(*t*), people infected and can infect new. People immune either by vaccination or by recovery from infection will be denoted by *R*(*t*). Finally deaths will be *M*(*t*). See [25] and [26] and the references therein for the use of mathematical models in epidemics.

We will model a period of 7 years. We will assume that population is constant. Our hypothesis establishes that the number of deaths for causes due to diphtheria is the same as the number of new borns. Mathematically it means that for every time *t*, the total number of population is a constant *N*, i.e. *S*(*t*) + *I*(*t*) + *R*(*t*) + *M*(*t*) = N.

The number of new infected by unit of time will depend on the number of encounters between infected and susceptible. Although only a proportion of these encounters will produce new infected. This proportion, that some times is called the “strength of the infection”, is denoted in our model by *β*(*t*). This parameter can change along the time, for example by the implementation of hygienic/prevention measures, such as household insulation of infected people, avoiding overcrowding, etc. The product of *β*(*t*) times *S*(*t*) and *I*(*t*), will be the new number of infected people who at the same time come from the susceptible class.

It is known that an immune individual can be carrier of the bacteria ([15]). In our model we don't contemplate this fact and we assume that the phenomenon is considered in the strength of infection *β*(*t*).

The amount by unit of time of infected people who leaves the class and goes either to the immune class or to the death class is, in theory, the proportion gives by inverse of the time the infection lasts. We shall call it γ.

In the case of the Russia epidemic we take *γ* = 49. This is because we use one year as a unit of time, and as the infection period lasts around a week, we adjust our model considering 49 weeks for a year. As the fatality rate is around 7%, this will give the proportion of infected people which goes to the class *M*(*t*).

The above thoughts are collected by the next system of differential equations

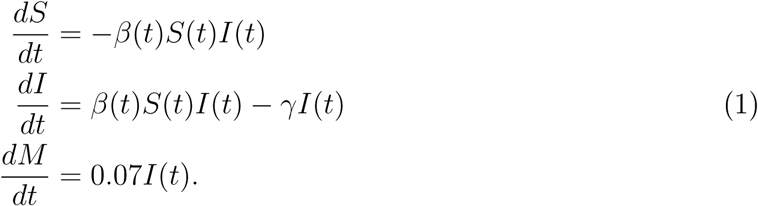

In order to solve it we need the initial value of the classes *S*(*t*)*, I*(*t*)*, R*(*t*) and *M*(*t*). The initial time *t*= 0 is for us the starting point of the soviet epidemic. The biggest cohort in Table 1 corresponds to Russia, we use this cohort as the main representative of the epidemic. Hence we choose an scenario with a total number of 100000 people and initial values *S*(0) = 35000, (we assume a vaccination of the 65%), *I*(0) = 0.82 (see Table 1) and *M*(0) = 0.

**Table 1.** Diptheria incidence in the Newly Independent States (NIS) and the Baltic States of the former Soviet Union, 1990-1998.

| Country | Population, in millions, 1994<sup>a</sup> | No. of cases/100,000 population |  |  |  |  |  |  |  |  |
| --- | --- | --- | --- | --- | --- | --- | --- | --- | --- | --- |
|  |  | 1990 | 1991 | 1992 | 1993 | 1994 | 1995 | 1996 | 1997 | 1998 |
| Western NIS and Russia |  |  |  |  |  |  |  |  |  |  |
| Russia | 147.37 | 1211 (0.82) | 1876 (1.27) | 3897 (2.64) | 15209 (10.32) | 39582 (26.86) | 35652 (24.19) | 13604 (9.23) | 4057 (2.75) | 1436 (0.97) |
| Belarus | 10.16 | 22 (0.22) | 26 (0.26) | 66 (0.65) | 120 (1.18) | 230 (2.26) | 322 (3.17) | 179 (1.76) | 102 (1.00) | 22 (0.22) |
| Ukraine | 51.47 | 109 (0.21) | 1103 (2.14) | 1553 (3.02) | 2982 (5.79) | 2990 (5.81) | 5280 (10.26) | 3156 (6.13) | 1364 (2.65) | 690 (1.34) |
| Subtotal | 209.00 | 1342 (0.64) | 3005 (1.44) | 5516 (2.64) | 18311 (8.76) | 42802 (20.48) | 41254 (19.74) | 16939 (8.10) | 5523 (2.64) | 2148 (1.03) |
| Baltic States |  |  |  |  |  |  |  |  |  |  |
| Estonia | 1.54 | 0 (0.00) | 7 (0.45) | 3 (0.19) | 11 (0.71) | 7 (0.45) | 19 (1.23) | 14 (0.91) | 3 (0.19) | 0 (0.00) |
| Latvia | 2.58 | 3 (0.12) | 5 (0.19) | 8 (0.31) | 12 (0.47) | 250 (9.69) | 369 (14.30) | 112 (4.34) | 42 (1.63) | 67 (2.60) |
| Lithuania | 3.71 | 2 (0.05) | 1 (0.03) | 9 (0.24) | 8 (0.22) | 38 (1.02) | 43 (1.16) | 11 (0.30) | 2 (0.05) | 2 (0.05) |
| Subtotal | 7.83 | 5 (0.06) | 13 (0.17) | 20 (0.26) | 31 (0.40) | 295 (3.77) | 431 (5.50) | 137 (1.75) | 47 (0.60) | 69 (0.88) |
| Caucasus and Moldova |  |  |  |  |  |  |  |  |  |  |
| Armenia | 3.55 | 7 (0.20) | 0 (0.00) | 0 (0.00) | 1 (0.03) | 36 (1.01) | 29 (0.82) | 11 (0.31) | 10 (0.28) | 4 (0.11) |
| Azerbaijan | 7.47 | 4 (0.05) | 66 (0.88) | 72 (0.96) | 160 (2.14) | 841 (11.26) | 883 (11.82) | 114 (1.53) | 31 (0.41) | 17 (0.23) |
| Georgia | 5.45 | 11 (0.20) | 7 (0.13) | 3 (0.06) | 28 (0.51) | 294 (5.39) | 419 (7.69) | 346 (6.35) | 288 (5.28) | 48 (0.88) |
| Moldova | 4.42 | 6 (0.14) | 14 (0.32) | 22 (0.50) | 35 (0.79) | 376 (8.51) | 418 (9.46) | 97 (2.19) | 49 (1.11) | 14 (0.32) |
| Subtotal | 20.89 | 28 (0.13) | 87 (0.42) | 97 (0.46) | 224 (1.07) | 1547 (7.41) | 1749 (8.37) | 568 (2.72) | 378 (1.81) | 83 (0.40) |
| Central Asian Republics and Kazakhstan |  |  |  |  |  |  |  |  |  |  |
| Kazakhstan | 17.03 | 28 (0.16) | 30 (0.18) | 45 (0.26) | 82 (0.48) | 489 (2.87) | 1106 (6.49) | 455 (2.67) | 162 (0.95) | 79 (0.46) |
| Kyrgyzstan | 4.67 | 6 (0.13) | 10 (0.21) | 4 (0.09) | 18 (0.39) | 304 (6.51) | 704 (15.07) | 412 (8.82) | 291 (6.23) | 138 (2.96) |
| Tajikistan | 5.93 | 11 (0.19) | 5 (0.08) | 16 (0.27) | 678 (11.43) | 1907 (32.16) | 4455 (75.13) | 1464 (24.69) | 723 (12.19) | 158 (2.66) |
| Turkmenistan | 4.01 | 4 (0.10) | 4 (0.10) | 22 (0.55) | 3 (0.07) | 43 (1.07) | 87 (2.17) | 80 (2.00) | 38 (0.95) | 19 (0.47) |
| Uzbekistan | 22.35 | 12 (0.05) | 13 (0.06) | 29 (0.13) | 137 (0.61) | 232 (1.04) | 639 (2.86) | 160 (0.72) | 34 (0.15) | 26 (0.12) |
| Subtotal | 53.99 | 61 (0.11) | 62 (0.11) | 116 (0.21) | 918 (1.70) | 2975 (5.51) | 6991 (12.95) | 2571 (4.76) | 1248 (2.31) | 420 (0.78) |
| Total NIS and Baltics | 291.71 | 1436 (0.49) | 3167 (1.09) | 5749 (1.97) | 19484 (6.68) | 47619 (16.32) | 50425 (17.29) | 20215 (6.93) | 7196 (2.47) | 2720 (0.93) |

NOTE. Data are no. of cases (incidence/100,000 population).
<sup>a</sup>Mid-year population (data source: United Nations Population Division).

The system above is solved numerically by using the software Sagemath. The Fourth Order Runge-Kutta method. In order to have a model that describes the Russia epidemic in Table 1, we found that the function *β*(*t*) could be given by the following graph.

**Figure 1.**
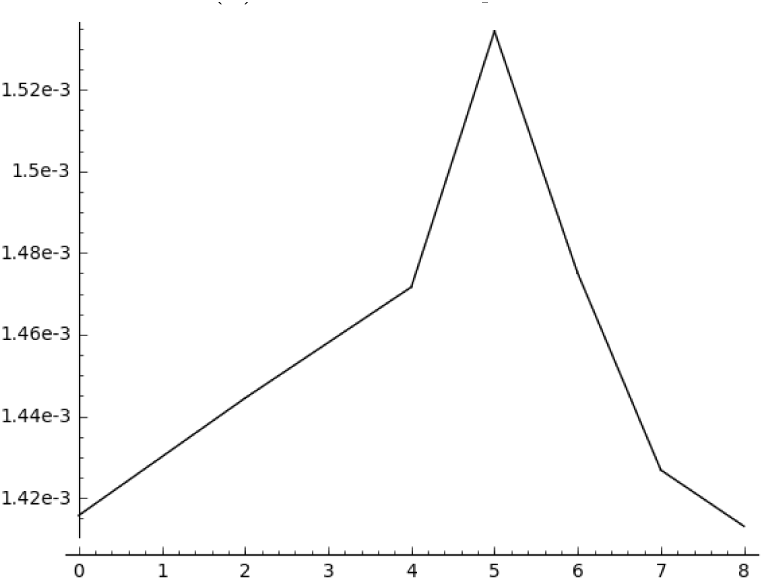
Graph of *β*(*t*)

The coincidence of our simulation with the real epidemic cases in Russia is given by the following figure containing the corresponding diagram and table.

**Figure 2.**
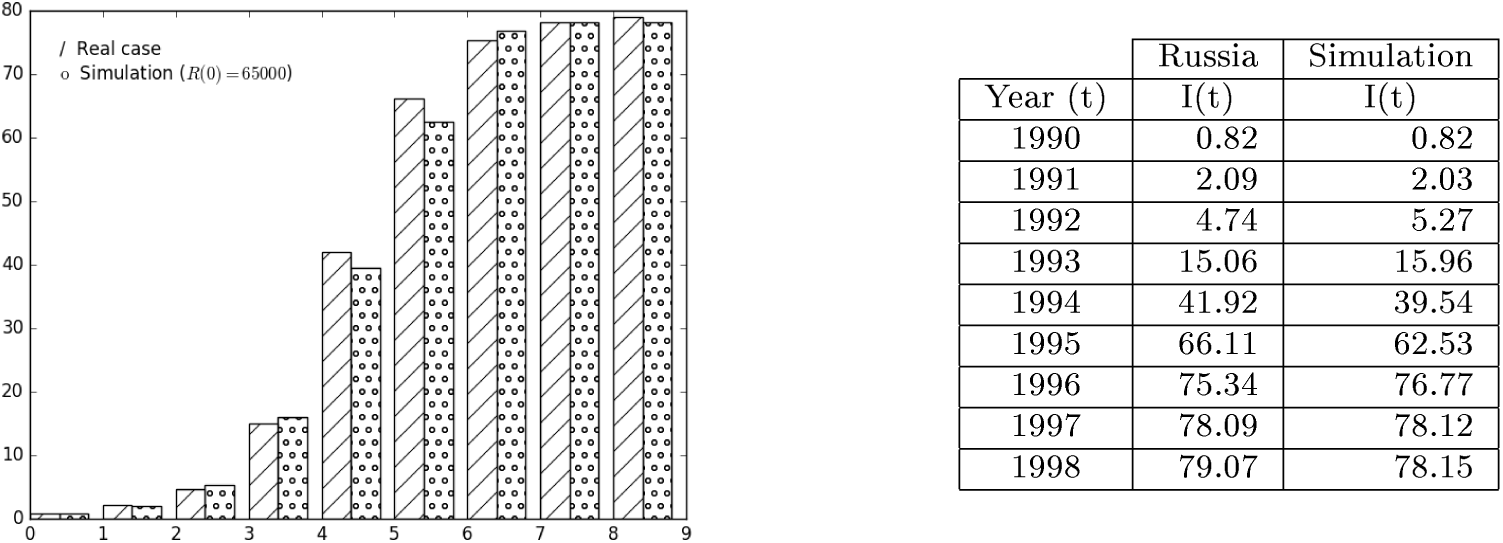
Simulation *R*(0) = 65000, *γ* = 49

## 3 Results

Once our mathematical model is adjusted to the data of the epidemic in Russia, we can simulate different scenarios.

### 3.1 Coverage vaccination goes down. Only 50% of population is covered.

Our first scenario analyzes the evolution of the epidemic under the hypothesis of a descent to 50% in vaccination coverage (it means 15% less coverage, that represents 23% if we take 65% as 100% coverage). If we decrease 23% the number of vaccinated people, we can see an increase of 47% in infected people. See Figure 3 contained the diagram and the corresponding table.

**Figure 3.**
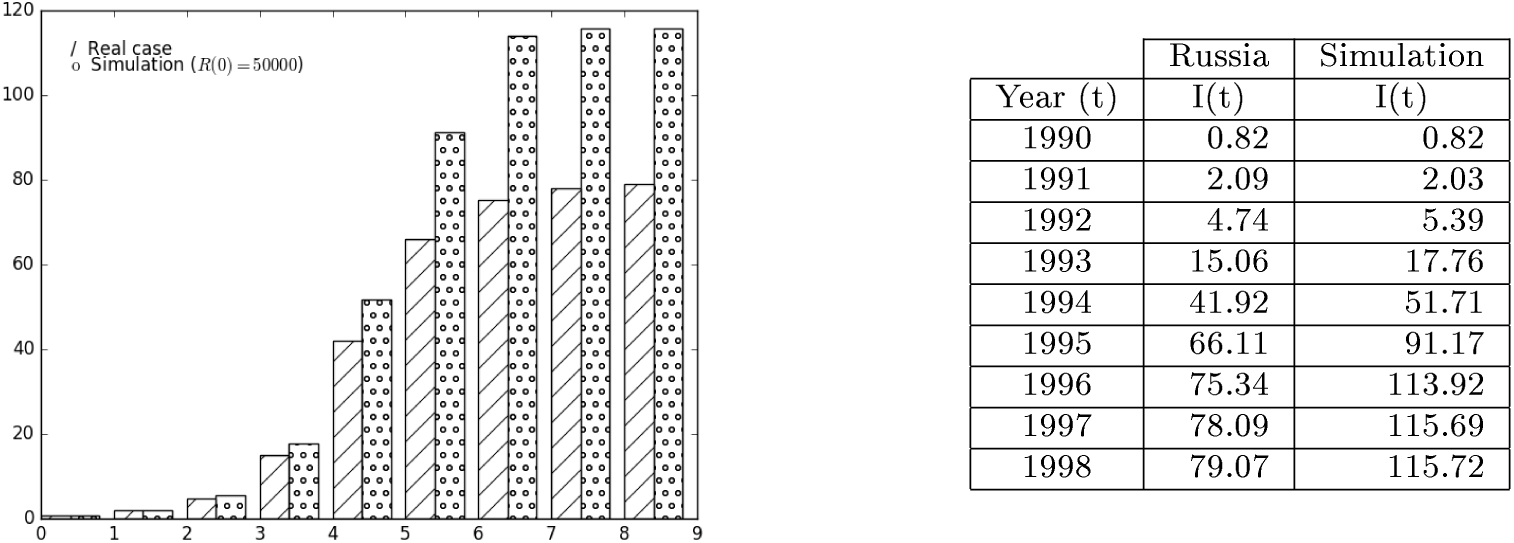
Simulation *R*(0) = 50000, *γ* = 49

### 3.2 Coverage vaccination goes up 80% or 90% of population

Opposite to this, we model the epidemic for the scenarios in which vaccine coverage rises either to 80% or to 90% (see Figure 4). We observe the herd immunity effect. A 80In the same order of ideas a 90

**Figure 4.**
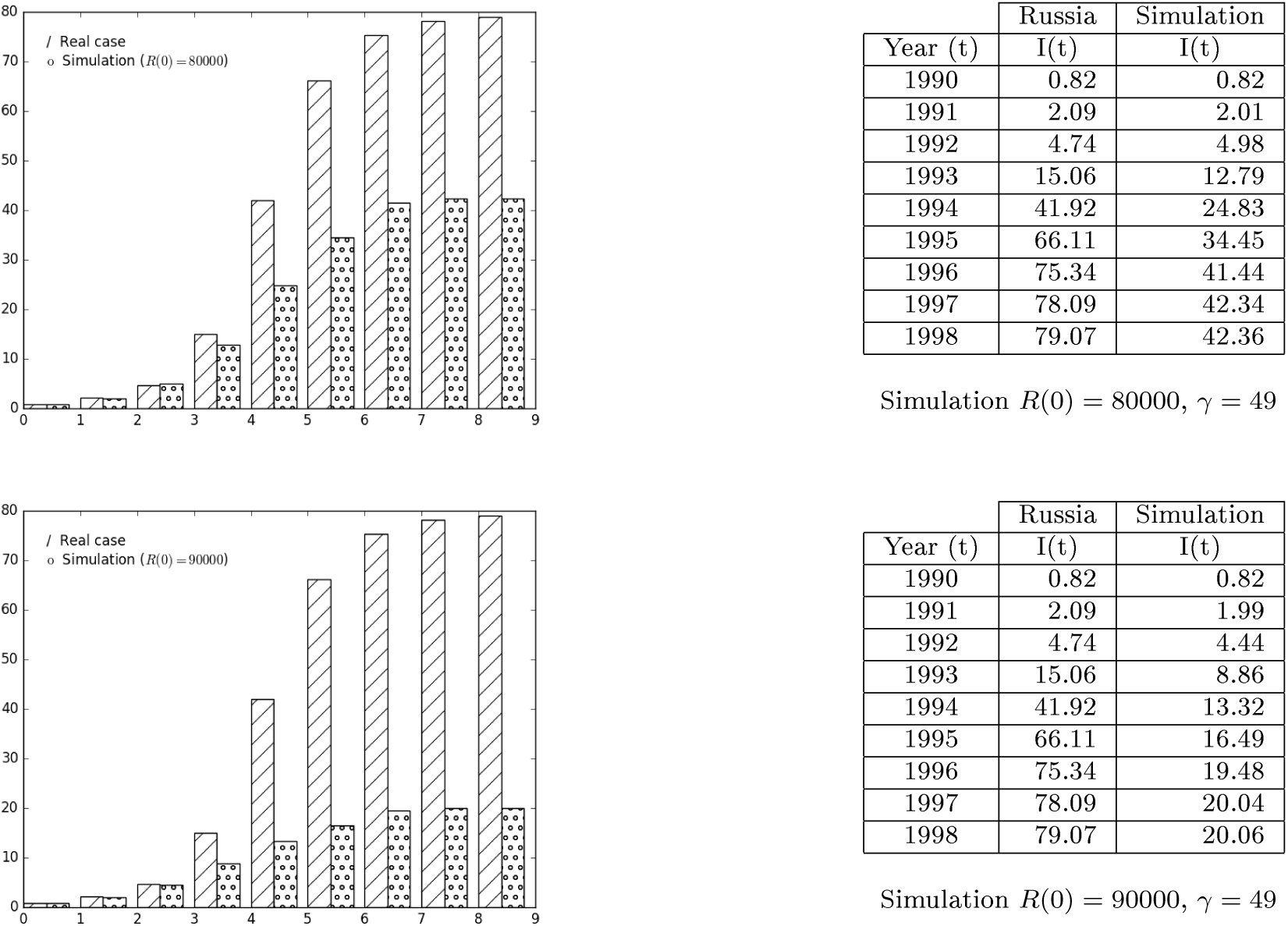
Increases of 23% and 38% in the vaccine coverage.

### 3.3 Public health care increases 5%.

As we have said before the epidemic can be disastrous in presence of poor health and sanitary conditions. As far as we know these conditions are present in refugee camps. In our model the parameter *γ* can be consider a measure of health care measures applied. Going from medicines, diphtheria antitoxine, appropriate toilets and tents to live and isolation of infected if necessary. We model a situation in which these conditions are implemented in such a way that the time of permanence of infected people in infection period is reduced in 5%. The numbers show that the amount of total infected people is reduced in a 11.31%. See Figure 5.

**Figure 5.**
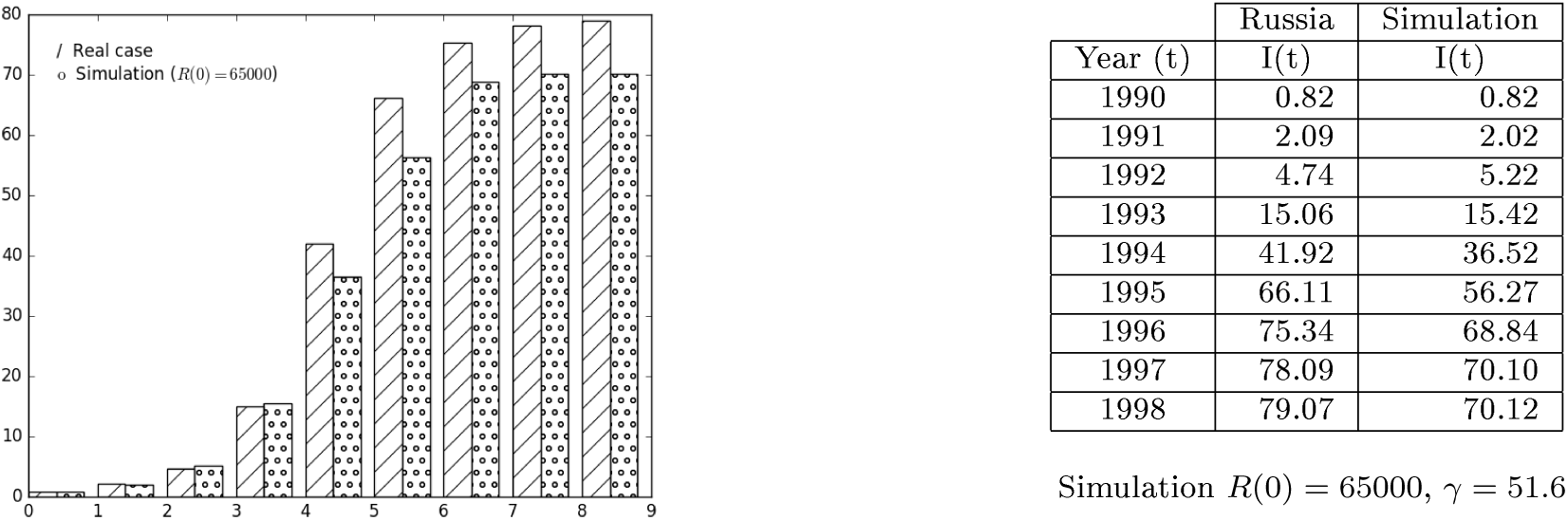
Increase of public health measures.

### 3.4 Combination of increasing health care and vaccination coverture.

Our model also can quantify a possible improvement of health measures together with an increase of vaccine coverage. We present in Figure 6 the diagram and table obtained in the case of an improvement of health measures in a 5% and a vaccine coverage of 75%. We observe that the reduction in this case rises to 38.6%.

**Figure 6.**
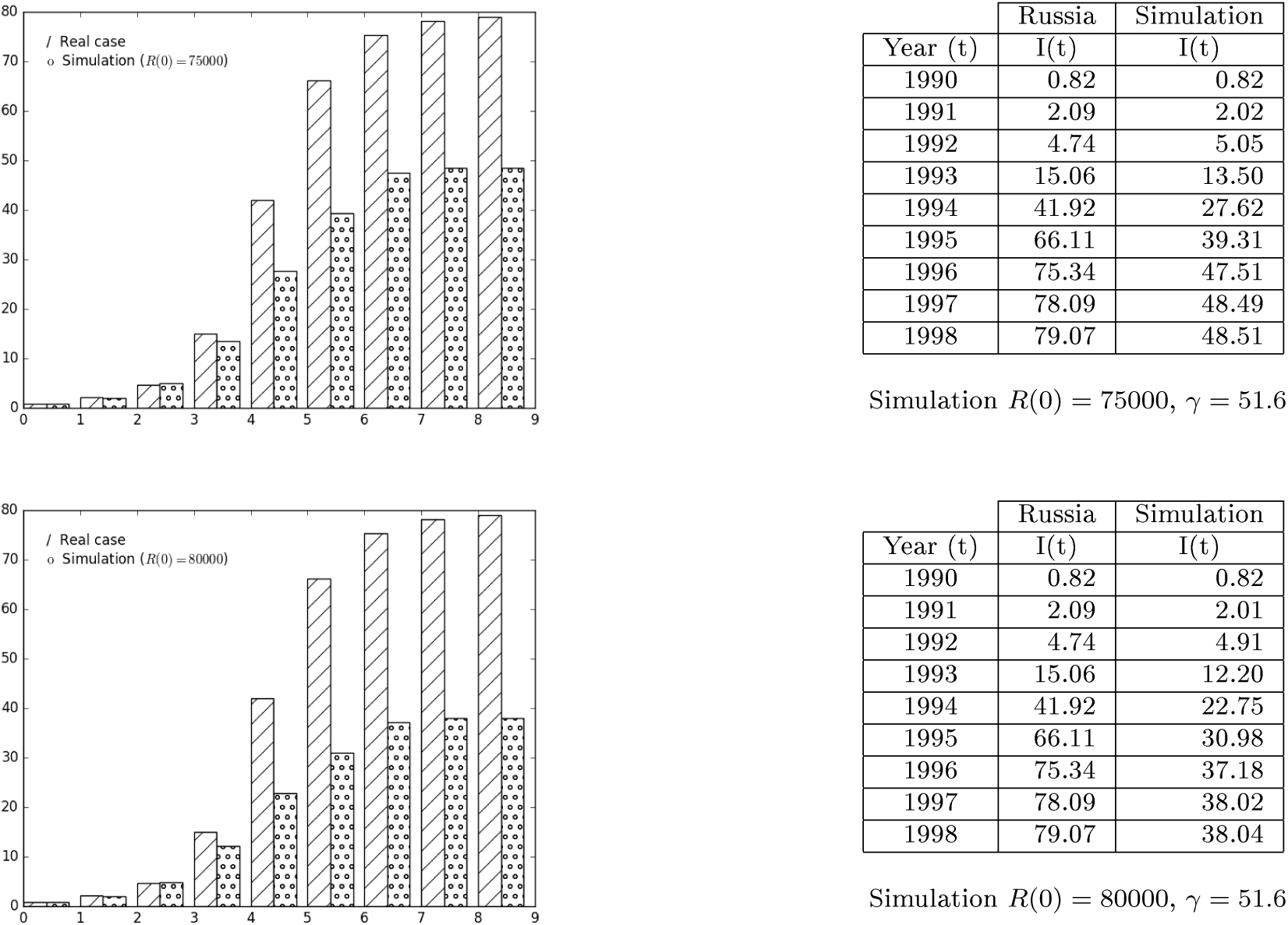
Increases of vaccine coverage and public health measures.

**Figure 7.**
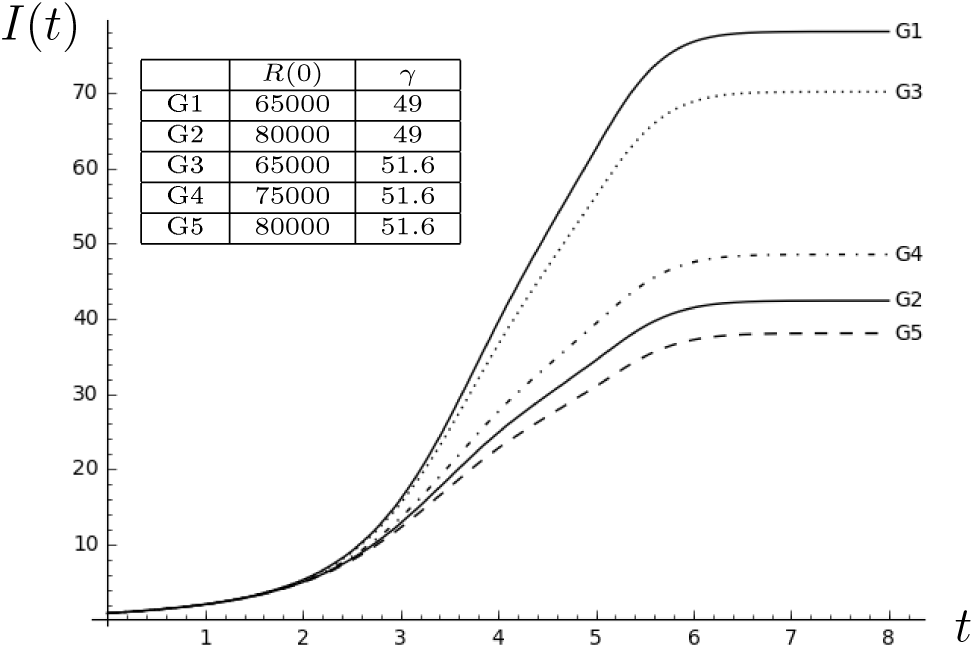
Comparison of different vaccine/health measures strategies.

Figure 6 shows the output of an improvement of health measures in 5% and a vaccine coverage of 80%. The reduction reaches to 51.89%.

## 4 Conclusions

Diphtheria was a very feared disease because of its high mortality rate. Luckily nowadays is a preventable disease thanks to its vaccine discovery. However, it has caused several epidemics before vaccination. One big epidemic occurred in the Soviet Union in 1990. Groups of experts coincide that the main causes of this outbreak were the reduction of vaccination coverage to 65% together with(see [2])

1. Large-scale population movements.
2. Socioeconomic instability.
3. Deterioration of health infrastructure.
4. Delay in implementing aggressive control measures
5. Lack of sources for prevention and treatment.

Our hypothesis of work is that this situation is being reproduced lately in movements of migrants in the Mediterranean area, mainly in the camps of Syrian refugees. Hence a new epidemic could start affecting mainly refugees but also could spread along the UE. In fact some cases have been already reported in Sweden, Germany and Denmark, see [10]. Our model deals with a temporal horizon of 7 years. We believe that diphtheria vaccine coverage among the refugees will not be able to reach the minimum desired rate of 90%, loosing the so called *herd innmnity efect* that describes how the ratio of protected people in a epidemic of direct transmission is bigger than the ratio of the vaccinated people, see [27].

Our model tries to quantify the effect of an improvement in public health conditions (given by different values of *γ*, see the system (1) and the comments about *γ* just before the system) together with some scenarios of vaccine coverage (given by the initial value *R*(0), which measures the vaccine coverage). Taking as reference a scenario G1 with a coverage of 65% and no public health measures, compared with 4 more possible scenarios we observed that:

- Compared with G2, 80% vaccine coverage: infected people are highly reduced
- With respect to G3, improving only health measures: the amount of infected people is greatly decreased
- In the case of G4, rising coverage to 75% and improving health measures: the number of infected people ameliorates, although not as much as with a higher vaccine coverage. We can see here the strength of the herd immunity effect
- Compared to the last scenario G5, with 80% of coverage and improving health measures: the proportion of infected population is significantly decrease. The graph refiects how much small increases in vaccine coverage and improvements in public health care can modified the disease evolution.

These estimations can be helpful in situations of social and economic instability, in order to prevent and control a disease spread.

